# Genetic Inhibition of Serum Glucocorticoid Kinase 1 Prevents Obesity-related Atrial Fibrillation

**DOI:** 10.1101/2021.05.20.444790

**Authors:** Aneesh Bapat, Guoping Li, Ling Xiao, Maarten Hulsmans, Maximillian J Schloss, Yoshiko Iwamoto, Justin Tedeschi, Xinyu Yang, Matthias Nahrendorf, Anthony Rosenzweig, Patrick Ellinor, Saumya Das, David Milan

**Author notes:** SD and DM contributed equally to this manuscript and share authorship. **Corresponding Author:** David Milan, MD, 265 Franklin St Ste 1902, Boston, MA 02110, E.

## Abstract

**Rationale:** Given its rising prevalence in both the adult and pediatric populations, obesity has become an increasingly important risk factor in the development of atrial fibrillation. However, a better mechanistic understanding of obesity-related atrial fibrillation is required. Serum glucocorticoid kinase 1 (SGK1) is a kinase positioned downstream of multiple obesity-related pathways, and prior work has shown a pathologic role for SGK1 signaling in ventricular remodeling and arrhythmias.

**Objective:** To determine the mechanistic basis of obesity associated atrial fibrillation and explore the therapeutic potential of targeting SGK1 in this context.

**Methods and Results:** We utilized a mouse model of diet induced obesity to determine the atrial electrophysiologic effects of obesity using electrophysiologic studies, optical mapping, and biochemical analyses. In C57BL/6J mice fed a high fat diet, there was upregulation of SGK1 signaling along with an increase in AF inducibility determined at electrophysiology (EP) study. These changes were associated with an increase in fibrotic and inflammatory signaling. Transgenic mice expressing a cardiac specific dominant negative SGK1 (SGK1 DN) were protected from obesity-related AF as well as the fibrotic and inflammatory consequences of AF. Finally, optical mapping demonstrated a shorter action potential duration and patch clamp revealed effects on *I*_*Na*_, with a decreased peak current as well as a depolarizing shift in activation/inactivation properties in atrial myocytes.

**Conclusions:** Diet induced obesity leads to increased cardiac SGK1 signaling as well as an increase in AF inducibility in obese mice. Genetic SGK1 inhibition reduced AF inducibility, and this effect may be mediated by effects on inflammation, fibrosis, and cellular electrophysiology.

## INTRODUCTION

The rate of obesity in the US has risen steadily since 1960, and reached a prevalence of 42% in 2017-2018.^1^ With a pediatric obesity prevalence of 19% in 2017-2018,^2^ there are no signs of slowing down. Atrial fibrillation (AF) is a common cardiac arrhythmia that affects about 5 million Americans and is associated with significant morbidity and mortality.^3^ Although many risk factors have been identified for AF, obesity has come to the forefront as a prominent-but potentially modifiable-one. Epidemiological studies demonstrate that nearly 1 in 5 cases of AF can be attributed to being overweight or obese.^4^ Importantly, data from the Women’s Health Study showed that obese individuals who lost weight to BMI<30 kg/m^2^ over a 5 year follow up period were found to have a reduced AF risk when compared to their persistently obese counterparts.^5^ Thus, not only is obesity a strong AF risk factor, but it may be one that is reversible.

The mechanistic relationship between obesity and AF is not completely delineated, and thus no specific therapies exist. However, a plethora of inciting factors have been proposed, and include alterations in hemodynamics, neurohormonal axis, inflammation, metabolism, and adipokines.^6^ Animal models of prolonged high fat diet feeding suggest a role for electroanatomic remodeling,^7–10^ inflammation,^11^ and decreased connexin expression.^12^ In addition, obesity is associated with activation of the NLRP3 inflammasome,^13^ which can be manipulated in cardiomyocytes to produce an atrial substrate susceptible to AF.^14^

Serum glucocorticoid kinase 1 (SGK1) is a PI3-kinase-dependent kinase, with structure similar to AKT.^15^ SGK1 expression and activation lies downstream of both insulin signaling pathways and mineralocorticoid receptor activation^16^-both of which are associated with human AF.^17–24^ Additionally, systemic upregulation of both upstream pathways have been noted in multiple models of diet induced obesity,^25–28^ positioning SGK1 as a potential key mediator of AF pathogenesis in the context of obesity. Initial studies of SGK1 demonstrated that the signaling pathway can be activated in the heart by transverse aortic constriction, where acute activation promotes cardiomyocyte survival.^29^ However, SGK1 was subsequently shown to be activated in human heart failure, pointing to its role in maladaptive cardiac remodeling. Cardiac specific expression of constitutively active SGK1 in mice was associated with worsened TAC-induced cardiac dysfunction, adverse electrical remodeling and lethal ventricular arrhythmias, while genetic inhibition was protective.^30^ SGK1 signaling has also been implicated in murine cardiac inflammation^31^ and NLRP3 inflammasome activation^32^ resulting from angiotensin II infusion induced hypertension. SGK1 signaling activates multiple pathways that overlap with the changes seen with obesity and metabolic syndrome, and hence presents an attractive potential therapeutic target.

We hypothesized that genetic inhibition of SGK1 would be protective in obesity-related AF through attenuation of obesity-related atrial electroanatomic remodeling and inflammation.

## METHODS

### Animal Studies

All studies were approved by the Institutional Animal Care and Use Committee (IACUC) and were in accordance with the NIH Guide for the Care and Use of Laboratory Animals. SGK1 dominant negative (DN) and constitutively active (CA) mice were generated as previously described, with cardiac-specific expression of a constitutively active (S422D) or dominant negative (K127M) SGK1 transgene driven by the α-myosin heavy chain promotor.^30^ Cardiac specificity was confirmed by immunoblotting for an incorporated hemagglutinin (HA) epitope tag. (Supplemental Figure 1A) Transgene carrying mice were bred with wild type C57BL/6J mice to generate litters of transgenic and WT mice. Mice were genotyped by PCR for the presence of the transgene using tail DNA and the following primers: forward 5’-GGTAGCAATCCTCATCGCTTTC-3’, reverse 5’-CTTCAGGGTGTTTGCATGCA-3’. (Supplemental Figure 1B) Starting at 6 weeks of age, the SGK1 DN and WT littermates were fed a high fat diet (Research Diets D12492, 60% fat by calories) for at least 10 weeks and up to 14 weeks until body weight was at least 34g. The weight cut-off was approximately the 95^th^ weight percentile among mice fed a control diet, and was used to exclude mice that may be resistant to diet induced obesity.^33^ Lean WT mice and SGK1 constitutively active mice were fed a control diet. Electrophysiology (EP) studies were performed under general anesthesia and terminated with extraction of the heart. Optical mapping and biochemical analyses required heart extraction, which was performed under deep anesthesia.

### Transthoracic Echocardiography

Transthoracic echocardiograms were performed in unanesthetized mice using a GE Healthcare Vivid E90 system equipped with a 15MHz linear probe (L8-18i-D). Left ventricular function and dimensions were determined using M-mode in a parasternal short axis view at the level of the papillary muscles. Left ventricular end diastolic diameter (LVEDD) and left ventricular end systolic diameter (LVESD) were measured at maximal and minimal diameters and fractional shortening calculated as the change in LV internal dimensions between end diastole and end systole normalized to the end-diastolic dimension [(LVEDD-LVESD)/LVEDD]. Left atrial size was measured as an anteroposterior dimension in a 2D parasternal long axis view. Gain was optimized to delineate the endocardial surfaces and measurements were made using the GE Healthcare EchoPAC workstation platform.

### Tail-Cuff Blood Pressure

Systolic, diastolic, and mean arterial pressures (SBP, DBP, MAP) were measured using a validated tail-cuff method that relies on volume pressure recording technology (Kent Scientific Corporation). Blood pressure measurements were made by using ten acclimation cycles followed by fifteen measurement cycles; the latter were averaged.

### Electrophysiology Studies

EP studies were performed under general anesthesia induced by administering 5% isoflurane driven by an oxygen source into an induction chamber. Anesthesia was subsequently maintained with 1%–2% isoflurane in 95% O_2_. An octapolar catheter (EPR-800, Millar) was inserted into the right jugular vein and positioned in the right atrium and ventricle. Sinus node function was determined by measuring the sinus node recovery time (SNRT) following 30 seconds of pacing at three cycle lengths (120, 100 and 80 ms). The Wenckebach cycle length was determined with progressively faster atrial pacing rates. Atrial, ventricular, and AV nodal refractory periods were measured using programmed electrical stimulation with overdrive pacing trains at 100ms followed by single extra-stimuli. Retrograde (VA) conduction Wenckebach cycle length was measured by pacing at progressively faster ventricular pacing rates. Provocative testing for arrhythmia induction was performed with triple extra-stimuli (S1-S2-S3) as well as rapid pacing at gradually faster rates to a pacing cycle length of 10ms. Atrial fibrillation (AF) was defined as a rapid atrial rhythm with atrial rate greater than ventricular rate and irregular ventricular response (R-R intervals). The duration of AF was measured from the end of the pacing train to the end of the rapid atrial activity. Mice were considered inducible for AF if they had episodes >1s in duration.

### Optical Mapping

Isolation and perfusion of the heart was performed as previously described.^34,35^ Briefly, the mouse was anaesthetized using isoflurane, the heart excised, and perfused via an aortic cannula. The cannulated heart was perfused with a modified Tyrode solution (128.2 mM NaCl, 4.7 mM KCl, 1.19 mM NaH_2_PO4, 1.05 mM MgCl_2_, 1.3 mM CaCl_2_, 20.0 mM NaHCO_3_, 11.1 mM glucose; pH 7.35±0.05) using a Langendorff perfusion setup. Blebbistatin (10 mM, Tocris Bioscience) was used to arrest cardiac motion. The heart was stained for 30 min with a voltage sensitive dye (di-4-ANEPPs, 2 mmol/L in Dimethyl sulfoxide). Custom-made epicardial platinum electrodes and a Medtronic stimulator were used to pace the heart. Pacing was performed at 60-120ms cycle lengths at twice the capture threshold (4 ms square wave stimuli). A halogen light source (X-Cite 150W, filtered at 520+45 nm) was used to excite fluorescence. Emissions >610 nm were collected and focused onto an 80 × 80 CCD camera (RedShirt Imaging SMQ Camera and Macroscope IIA) using a 50 mm ×2.7 lens (numerical aperture 0.4). Data sampling was performed at 2000 frames per second with a filter setting of 1 kHz. A specifically designed Matlab program was used to perform data analysis in order to generate conduction velocities and APD at 50, 70, and 90% repolarization.

### Cardiomyocyte Isolation

Mice were anaesthetized as described above and intraperitoneally injected with 0.2 cc heparin. Hearts were then dissected and mounted on a Langendorff system (Radnoti) via aortic cannulation. Hearts were perfused with Ca^2+^-free normal Tyrode’s solution containing NaCl 137 mM, KCl 4 mM, MgCl_2_ 1mM, HEPES 10 mM, NaH_2_PO_4_ 0.33 mM, taurine 5mM and dextrose 10 mM (pH 7.4), followed by the same perfusion buffer containing collagenase D and B (Roche) and protease XIV (Sigma) at 37°C. After enzymatic digestion, LA appendage was dissected into Kraftbrühe (KB) buffer containing KCl 25 mM, KH_2_PO_4_ 10 mM, dextrose 20 mM, DL-aspartic acid potassium salt 10 mM, bovine albumin 0.1%, L-glutamic acid potassium salt 100 mM, MgSO_4_ 2 mM, taurine 20 mM, EGTA 0.5 mM, creatine 5 mM, HEPES 5 mM (pH 7.2). Single LA cardiomyocytes were dispersed by triturating the digested tissue and seeded on laminin-coated 8-mm coverslips. After 1-hour incubation at 37°C in 5% CO_2_ incubator, cardiomyocytes were treated with a gradually increased Ca^2+^ concentration from 0.06 mM to 1.2 mM in normal Tyrode’s solution at RT.

### Patch Clamping

Sodium currents (*I*_*Na*_) were recorded by whole-cell patch-clamp techniques in isolated murine LA cardiomyocytes at RT. Borosilicate-glass electrodes with tip-resistances 1.5-3 MΩ were filled with pipette solution. For I_*Na*_ recordings, pipette solution contained CsCl 135 mM, NaCl 10 mM, CaCl_2_ 2 mM, EGTA 5 mM, HEPES 10 mM, MgATP 5 mM (pH 7.2 with CsOH); bath solution contained NaCl 50 mM, CaCl_2_ 1.8 mM, MgCl2 1 mM, CsCl 110 mM, dextrose 10 mM, HEPES 10 mM, nifedipine 0.001 mM (pH 7.4 with CsOH). Currents were low-pass filtered at 5 kHz with an Axopatch 200B amplifier and digitized at 10 kHz with a Digidata 1440A A/D converter. I_*Na*_ was obtained with 50-ms depolarizations (−100 mV to + 40 mV) from a holding potential of - 120 mV at 0.1 Hz. Data were recorded with Clampex 10.3 and analyzed with Clampfit 10.3 (Molecular Devices Inc.). Cell-capacitance and series resistance were compensated by ~70%. Leakage compensation was not used. Currents were normalized to cell capacitance.

### Flow Cytometry

Mice were perfused through the left ventricle with 10 mL of ice-cold PBS. Hearts were excised and atrial tissue was micro-dissected using a dissection microscope. After harvest, tissues were minced into small pieces and subjected to enzymatic digestion with 450 U/mL collagenase I, 125 U/mL collagenase XI, 60 U/mL DNase I and 60 U/mL hyaluronidase (all Sigma-Aldrich) for 40 minutes at 37°C under agitation. Tissues were then triturated, and cells were filtered through a 40-µm nylon mesh, washed, and centrifuged to obtain single-cell suspensions. For a myeloid cell staining on processed heart samples, isolated cells were stained at 4°C in FACS buffer (PBS supplemented with 0.5% BSA) with mouse hematopoietic lineage markers including PE-conjugated anti-mouse antibodies directed against B220 (clone RA3-6B2, 1:600), CD49b (clone DX5, 1:1200), CD90.2 (clone 53–2.1, 1:3000), CD103 (clone 2E7, 1:600), Ly6G (clone 1A8, 1:600), NK1.1 (clone PK136, 1:600), and Ter-119 (clone TER-119, 1:600). This was followed by a second staining for CD11b (clone M1/70, 1:600), CD45 (clone 30-F11, 1:600), F4/80 (clone BM8, 1:600) and Ly6C (clone HK1.4, 1:600). DAPI was used as a cell viability marker. Neutrophils were identified as CD45highCD11bhigh(B220/CD49b/CD90.2/CD103/NK1.1/Ter-119)lowLy6Ghigh and cardiac macrophages as CD45highCD11bhigh(B220/CD49b/CD90.2/CD103/Ly6G/NK1.1/Ter-119)lowF4/80highLy6Clow/int. Antibodies were purchased from BioLegend and BD Biosciences. Data were acquired on an LSRII (BD Biosciences) and analyzed with FlowJo software.

### Quantitative PCR of RNA

Tissue samples were snap frozen and then lysed using a Tissue Lyser (Qiagen) in TRIzol (Thermo Fisher) followed by RNA extraction. A total of 500ng of RNA was reverse transcribed using the High Capacity cDNA Reverse Transcription Kit (Thermo Fisher). Real-time PCR was conducted with SsoAdvance Universal SYBR Green (Bio-Rad) using a QuantStudio 6 Flex Real-Time PCR System (Thermo Fisher). Quantification of all target-gene expression was normalized to β-actin. Primers for genes of interest are presented in supplemental table 1.

### Immunoblotting

Western blot analysis was performed on snap-frozen atrial tissue. Tissue samples were lysed using a Tissue Lyser (Qiagen) in protein lysis buffer supplemented with protease and phosphatase inhibitors (Boston BioProducts). Protein content was measured using a commercially available kit (DC Protein Assay, Bio-Rad). Equal amounts (15-25 μg) were treated with Laemmli buffer and β-mercaptoethanol and incubated at 100° C for 5 minutes. The lysate was then electrophoresed on a 4-20% SHS-polyacrylamide resolving gel and transferred to a PVDF membrane. The membrane was incubated overnight at 4° C with relevant antibodies, and then hybridization was completed with secondary antibody for 2 hours at room temperature. Antibody signal detection was achieved by employing the Clarity Western ECL Substrate (BioRad #1705061). Imaging and image quantification were done via BioRad Chemidoc Touch Imaging System and ImageLab, respectively.

### Histology and Fibrosis Quantification

Mice hearts were dissected and perfused with PBS. Hearts were then embedded in Tissue-Tek O.C.T. compound (Sakura Finetek), snap-frozen in 2-methylbutane on dry ice and sectioned into 10 μm slices using CryoJane Tape-Transfer System (Leica). Masson’s trichrome stain for cardiac fibrosis was performed according to the manufacturer’s instructions (Sigma). Quantification of fibrosis in the atria was performed using the BZX Analyzer software.

### Cytokine ELISAs

Cardiac puncture was performed during heart extraction to collect plasma. Commercially available kits were utilized to determine levels of IL-6 and CRP (R&D systems).

### Statistics

All statistical analyses were conducted with GraphPad Prism software. Animal group sizes were as low as possible and were empirically chosen. Categorical variables with binary outcomes were compared with the Fisher exact test. For two group comparisons of continuous variables, two-tailed unpaired or paired Student’s t-test was used to determine statistical significance. If a normal distribution could not be assumed, statistical significance was evaluated using the two-tailed Mann-Whitney test. P values < 0.05 were used to denote significance.

## RESULTS

### Diet induced obesity results in increased AF susceptibility and increased SGK1 signaling

We used diet induced obesity to create a mouse model of obesity-related AF. C57BL/6J wild type (WT) mice were fed a high fat diet (HFD) starting at 6 weeks of age. (Figure 1A) After 4 to 10 weeks of HFD, mice underwent terminal EP studies and tissues were harvested. Although none of the mice (0 out of 3) were inducible for AF greater than 1 second after 4 weeks of HFD (average weight 31±2.2 grams), 6 out of 7 were inducible for AF after 10 weeks of HFD (average weight 42±3.1 grams). In comparison, among age-matched mice fed a control diet for 10 weeks (average weight 30±2.8 grams), only 1 out of 8 was inducible for AF during EP studies. (Figure 1B) Given this stark phenotype at 10 weeks of HFD, we proceeded to use 10 weeks of HFD as a model for obesity-related AF.

**Figure 1:**
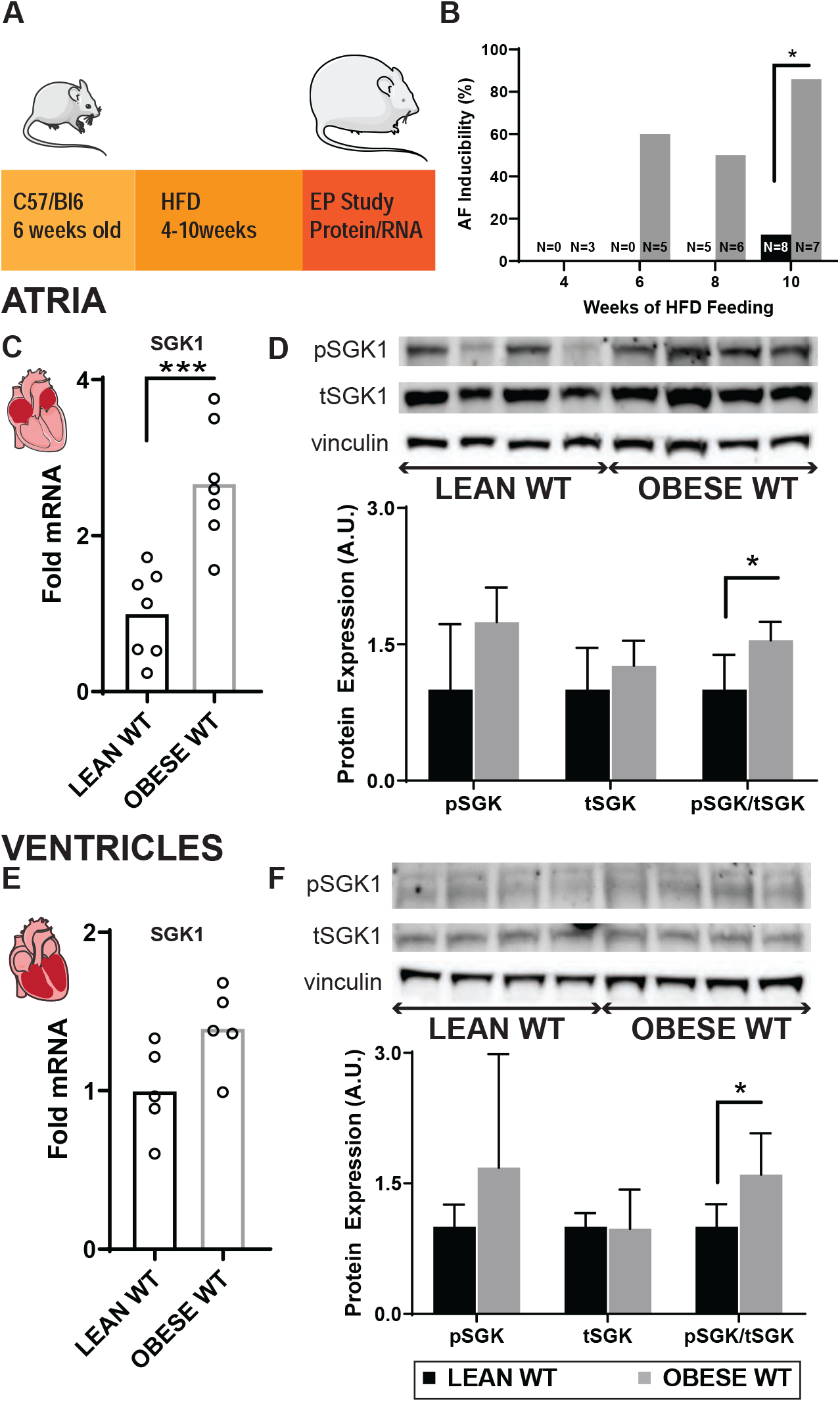
Diet induced obesity results in increased AF inducibility and is associated with upregulation of SGK1 signaling. *A*, Schema of mouse model using C57BL/6J mice. *B*, AF inducibility in mice fed a HFD (black) or control diet (grey) for the specified period of time. *C*, Atrial SGK1 mRNA expression in obese versus lean mice. *D*, Expression of atrial phosphorylated (pSGK1) and total (tSGK1) SGK1 protein as quantified by Western blotting. *E*, Ventricular SGK1 mRNA expression in obese versus lean mice. *F*, Expression of ventricular phosphorylated (pSGK1) and total (tSGK1) SGK1 protein as quantified by Western blotting.

In addition to EP studies, a separate group of WT mice were euthanized after 10 weeks HFD to harvest cardiac tissue and determine SGK1 activation. The tissue was used to extract and analyze protein and mRNA expression. HFD resulted in a significant increase in atrial SGK1 gene expression, with a similar trend in the ventricles. (Figure 1C,E) In addition, in both atrial and ventricular tissue, high fat diet feeding led to a significant increase in the ratio of phosphorylated to total SGK1 as determined by Western blotting. (Figure 1D, F) Overall, these data demonstrate that diet induced obesity results in increased AF inducibility and is associated with an increase in SGK1 transcription and activation.

### Genetic SGK1 inhibition alters atrial electrophysiology and prevents obesity-related AF

In order to test whether SGK1 plays a role in obesity-induced AF we employed a previously described transgenic mouse model that overexpresses a dominant negative form of SGK1 in cardiomyocytes (SGK1 DN).^30^ After 10-14 weeks of high fat diet, 79% of SGK1 DN (23 out of 29) and 73% of WT littermates (22 out of 30) achieved the minimum 34g weight. Over the course of feeding, there were no differences in weight gain or the final weight between SGK1 DN and WT littermates. Two-dimensional transthoracic echocardiography (TTE) was performed and did not reveal any differences in LV systolic or diastolic dimensions, fractional shortening or left atrial size between SGK1 DN and WT mice. Tail cuff blood pressure measurements showed no difference in systolic, diastolic, or mean arterial pressure. (Supplemental Figure 2)

Terminal EP studies were performed to determine EP parameters with focus on AF inducibility. Parameters related to sinus node function, AV node function, and refractory periods in atria and ventricles are presented in Table 1. There were no significant differences between the two groups at baseline. Programmed electrical stimulation with up to 2 extra-stimuli and rapid pacing were performed in both the atria and ventricles. There was a marked difference in AF inducibility between the two groups (Figure 2). While all (7 out of 7) WT mice were inducible for AF, genetic inhibition of SGK1 was significantly protective in terms of AF inducibility (2 out of 8). This finding was further supported by the observation that during the entirety of the EPS, even short (>250ms) episodes of AF were induced less frequently, and there was a lower total AF burden in SGK1 DN compared to WT mice.

**Table 1.**
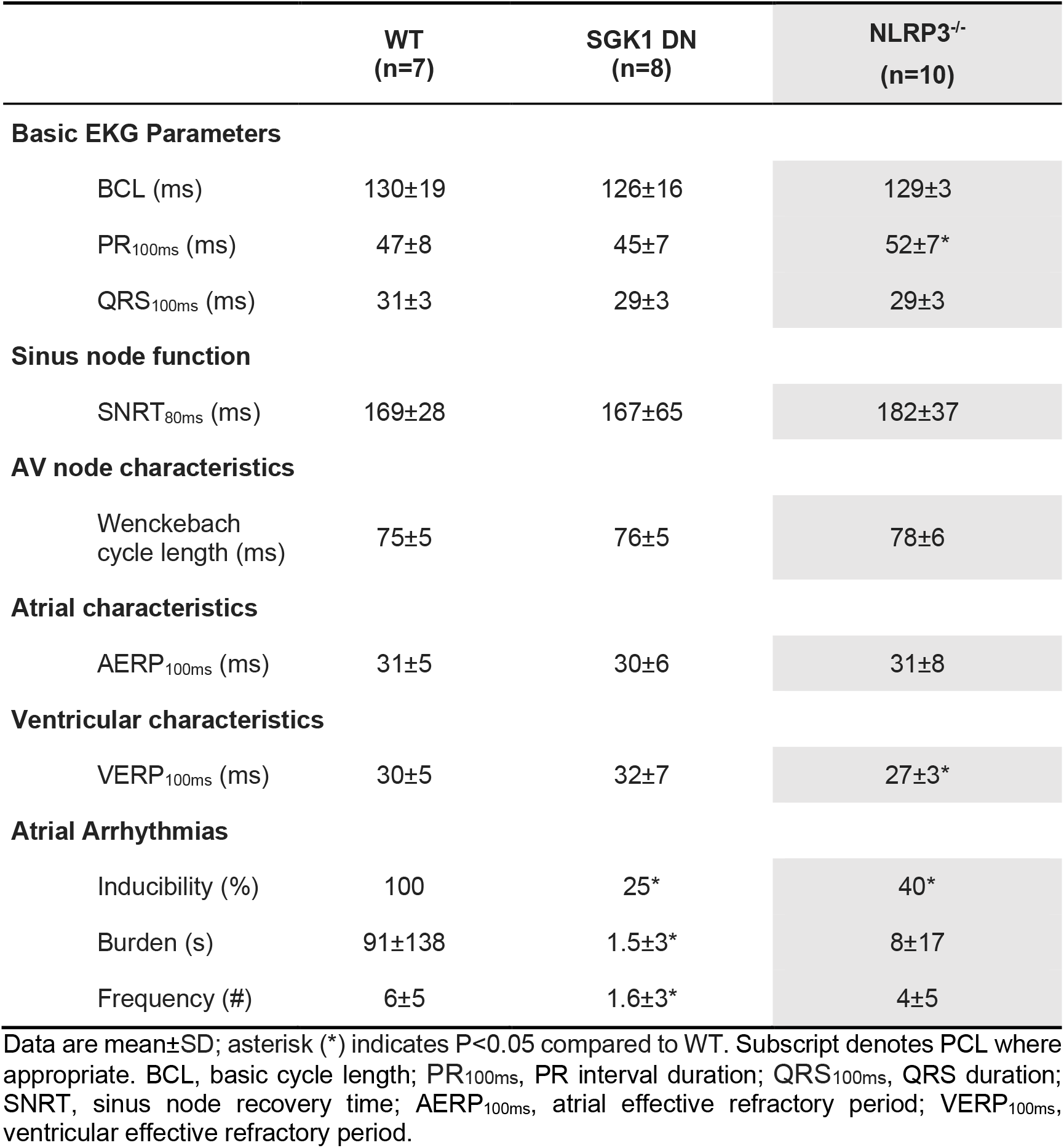
Cardiac electrophysiological parameters determined with in vivo EP study in obese WT, SGK1 DN, and NLRP^-/-^ mice.

**Figure 2:**
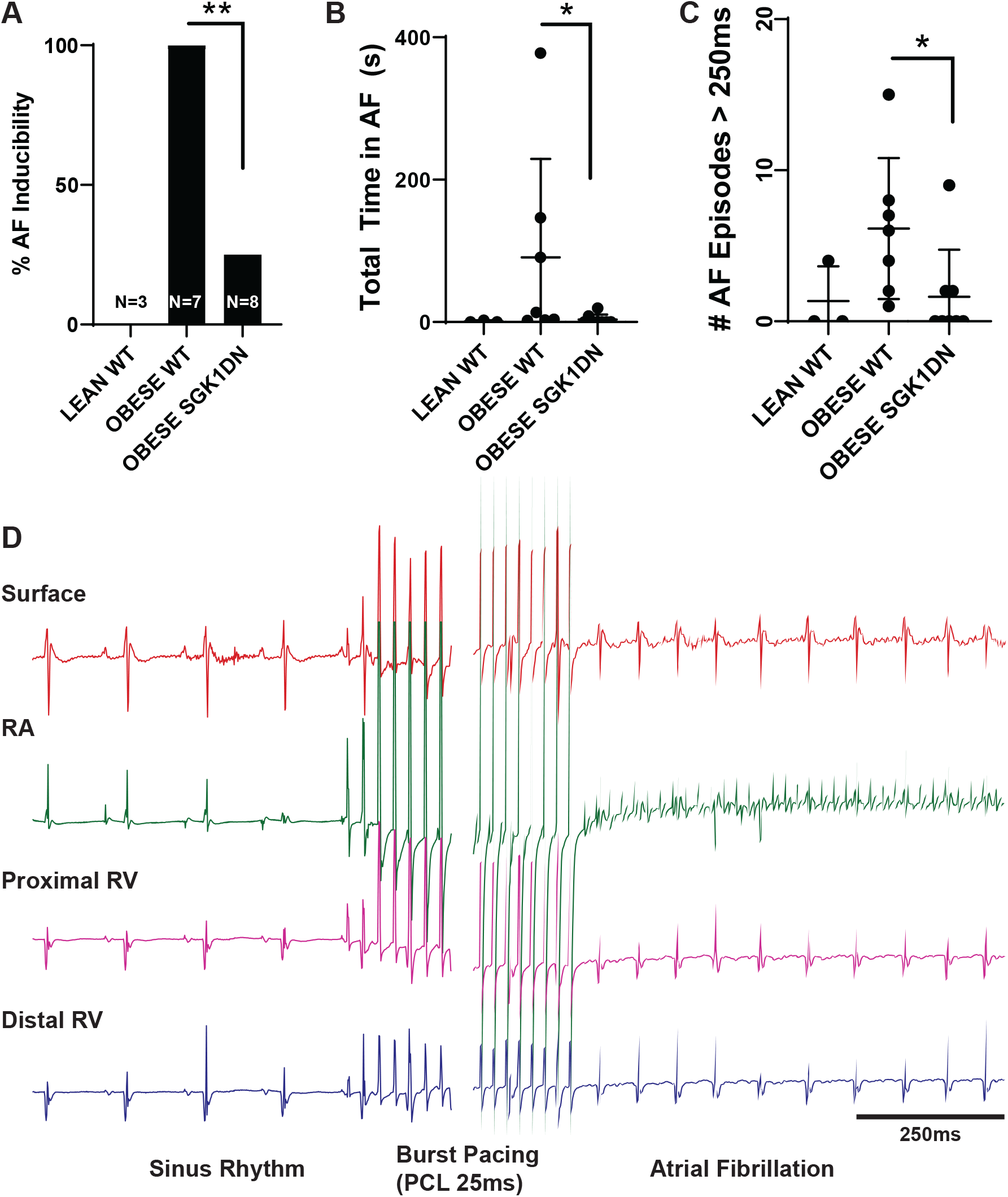
SGK1 DN mice are protected from obesity-induced AF. *A*, AF inducibility, by percent inducible, of lean and obese mice. *B*, Total summed duration of all AF episodes during each EP study. *C*, Number of episodes of induced AF > 250ms in duration by mouse. *D*, Example intracardiac tracing showing pacing-induced AF.

Optical mapping of right atrial and left atrial appendages was performed on SGK1 DN and WT mice fed HFD using a voltage sensitive dye. (Table 2) Overall, the action potential duration was generally shorter in the obese SGK1 DN as compared to the obese WT mice in both atrial chambers and met statistical significance at 90% repolarization (APD90). (Figure 3A) Additionally, there was a trend towards a decreased left atrial dV/dt in the SGK1 DN mice (WT 34±3.6%/ms; DN 31.4±4.3%/ms; p=0.07). There were no significant differences in conduction velocity (CV) between the two obese mouse genotypes. Interestingly, within the WT obese mice, there were significant inter-atrial differences in APD90, CV and dV/dt; SGK1 DN mice only had an analogous interatrial difference in CV. (Figure 3B) Overall these data suggest that SGK1 activity plays a role in defining atrial and interatrial electrophysiology.

**Table 2.** Optical mapping derived action potential durations, conduction velocity, and AP upstroke velocity in WT and SGK1 DN atria.

|  | WT | SGK1 DN | *P* value |
| --- | --- | --- | --- |
| **Right Atrium** | **N=15** | **N=19** |  |
| CV (m/s) | 0.67±0.26 | 0.61±0.14 | 0.42 |
| APD50 (ms) | 11.1±1.2 | 10.0±1.7 | **0.04** |
| APD70 (ms) | 17.9±2 | 16.1±2.6 | **0.04** |
| APD90 (ms) | 22±2.7 | 19.6±2.7 | **0.03** |
| dV/dt (%/ms) | 29±4.4 | 30.5±5.3 | 0.48 |
| **Left Atrium** | **N=11** | **N=15** |  |
| CV (m/s) | 0.78±0.31 | 0.74±0.22 | 0.68 |
| APD50 (ms) | 9.4±1.3 | 8.9±1.7 | 0.41 |
| APD70 (ms) | 15.9±2.7 | 14.6±2.6 | 0.21 |
| APD90 (ms) | 20.9±2.5 | 18.0±2.9 | **0.02** |
| dV/dt (%/ms) | 34±3.6 | 31.4±4.3 | 0.07 |

Data are mean±SD; unpaired *t* test. PCL 100ms for all measurements. CV, conduction velocity; APD50, action potential duration at 50% repolarization; APD90, action potential duration at 90% repolarization; dV/dt, AP upstroke velocity from 20-80% depolarization calculated as percent per ms.

**Table 3.**
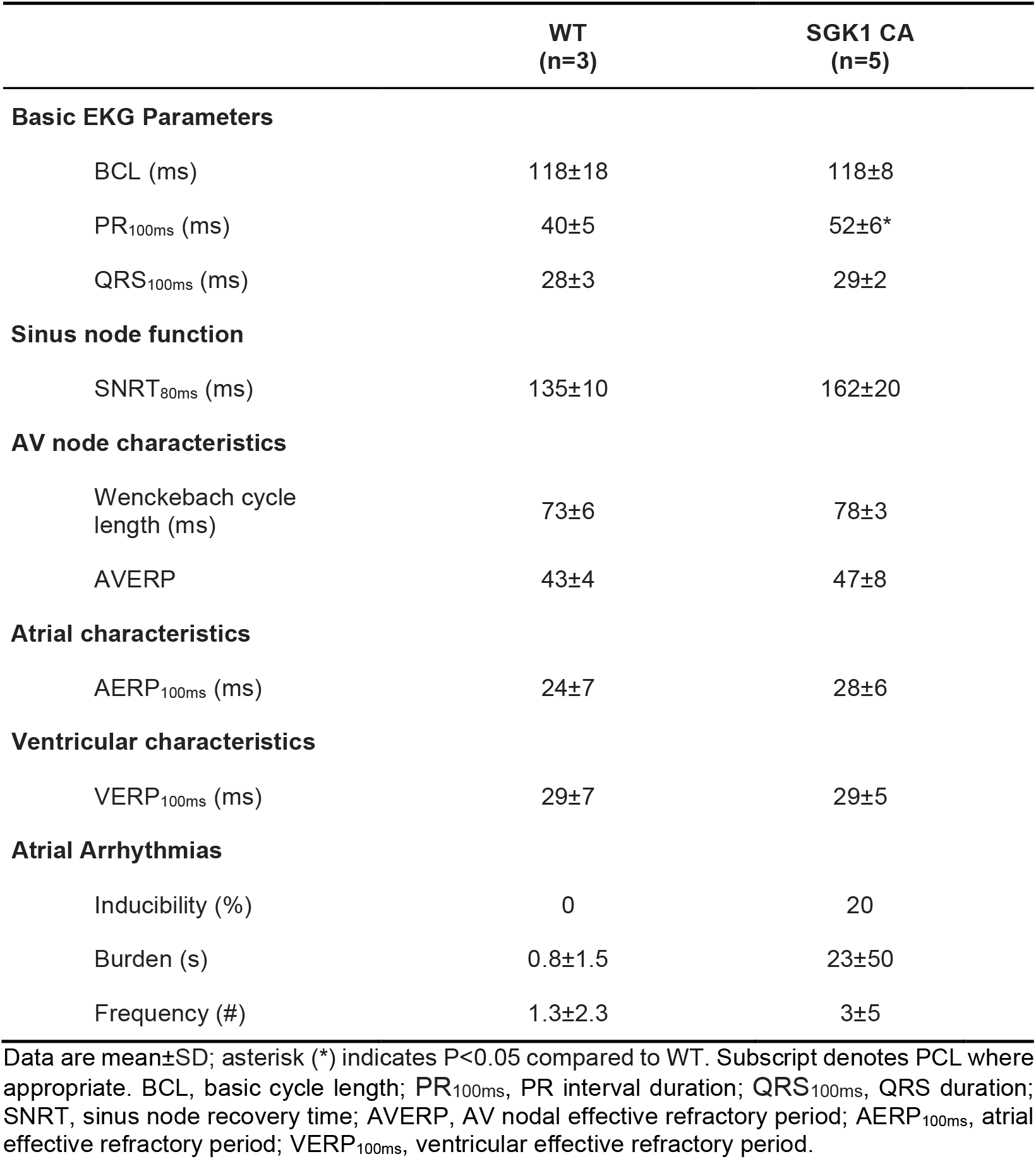
Cardiac electrophysiological parameters determined with in vivo EP study in lean WT and SGK1 CA mice.

**Figure 3:**
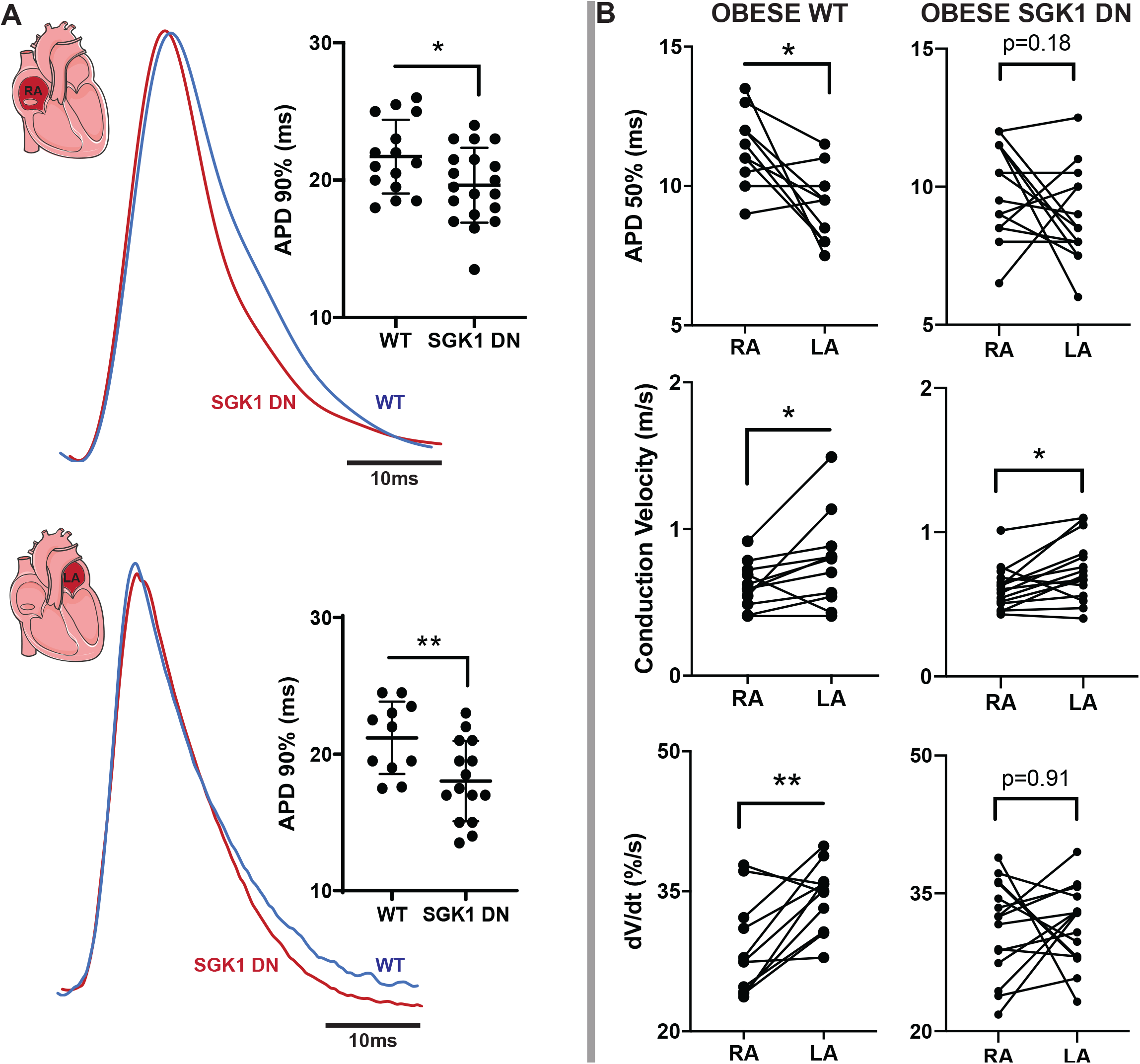
Atrial electrophysiologic effects of SGK1 genetic inhibition on obese mice. *A*, Right (top) and left (lower) atrial action potential durations at 90% repolarization in obese wild type and SGK1 DN mice. Tracings show representative optical action potential tracings. *B*, Interatrial differences in APD at 50% repolarization (top), conduction velocity (middle), and upstroke velocity (dV/dt, bottom) in obese WT (left) and SGK1 DN (right) mouse atrial chambers.

We performed focused patch clamping on atrial myocytes to further investigate the differences in APD and dV/dt found in optical mapping. Prior work has demonstrated the effect of SGK1 activation on sodium current via alteration of channel biophysical properties, as well as trafficking and localization of Na_V_1.5, the pore forming subunit of the channel.^30^ In addition, adipokines such as leptin have been shown to increase the late sodium current in left atrial myocytes.^36^ We suspected that the protective electrophysiologic consequences of SGK1 genetic inhibition may result from its effects on the sodium current. Whole cell patch clamp revealed that SGK1 DN left atrial myocytes-as compared to WT left atrial myocytes-had a significantly lower peak *I*_*Na*_ as well as a depolarizing shift in the steady state activation/inactivation properties. (Figure 4A-C) These data are consistent with previously published data regarding the effect of SGK1 activation/inactivation on *I*_*Na*_ in ventricular myocytes^30^ and may account for the trend toward reduced dV/dt in the SGK DN mice.

**Figure 4:**
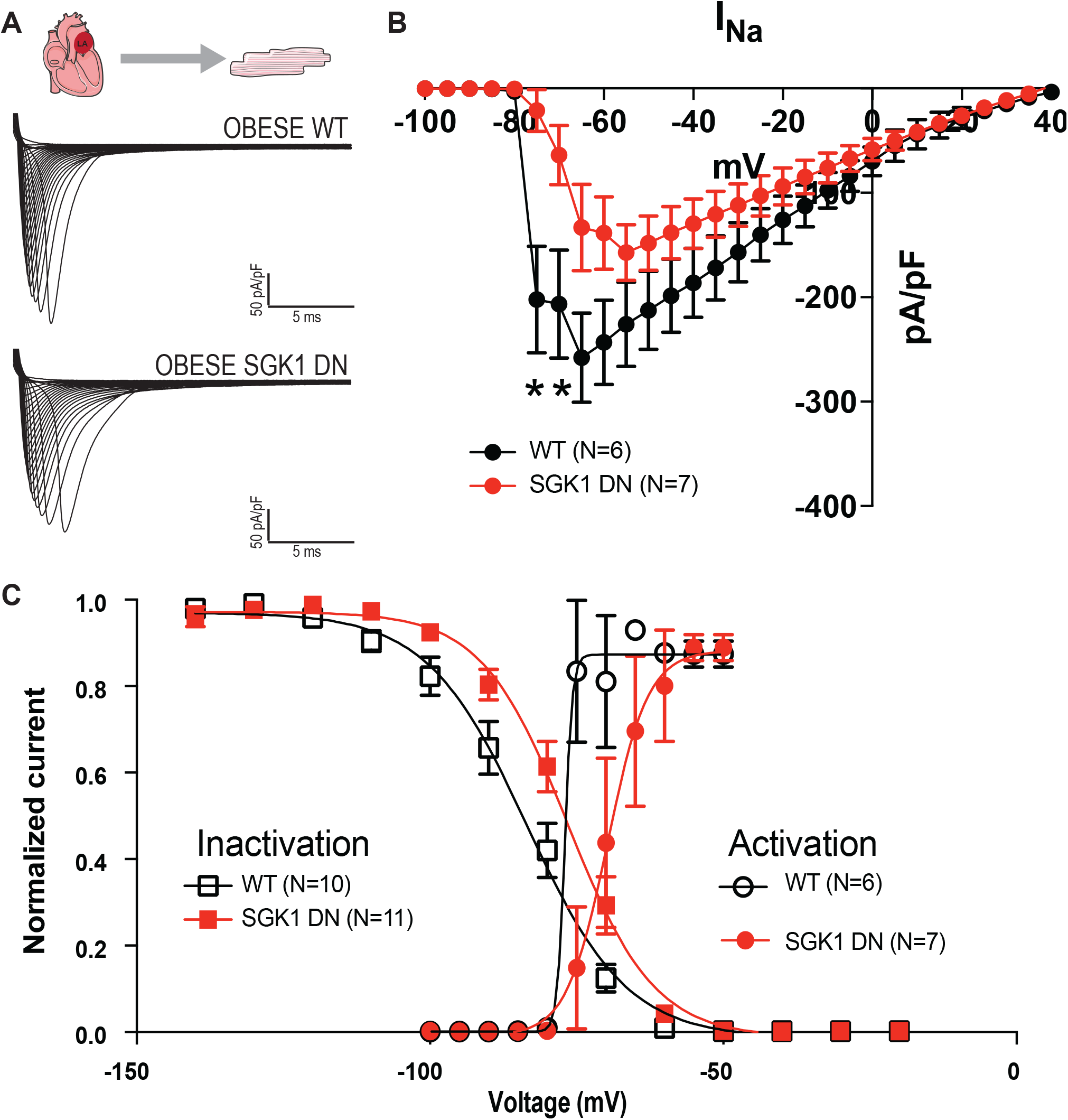
Effect of SGK1 genetic inhibition on *I*_*Na*_ in obese left atrial cardiomyocytes. *A*, Example sodium current recordings from isolated left atrial (LA) cardiomyocytes. *B*, Current-voltage relations in wild type (black) and SGK1 DN (red) isolated LA cardiomyocytes demonstrating a significant difference in peak *I*_*Na*_. *C, I*_*Na*_ activation-inactivation voltage dependences for WT (black) and SGK1 DN (red) left atrial cardiomyocytes demonstrating a depolarizing shift in the SGK1 DN myocytes.

### Genetic inhibition of SGK1 prevents obesity-induced cardiac remodeling

SGK1 DN mouse ventricles are resistant to TAC-induced fibrosis,^30^ so we hypothesized that SGK1 inhibition may protect from obesity-induced structural remodeling. Although there were no significant differences in fibrosis as measured by Masson-Trichrome staining between the three groups (Figure 5A), we suspected that this may be an insensitive technique. Western blotting revealed a marked increase in TGF-β protein expression in all obese – as compared to lean-mice, with no difference based on genotype. However, protein expression of connective tissue growth factor (CTGF) was reduced in SGK1 DN as compared to WT obese mouse atria. (Figure 5B) In addition, quantitative PCR demonstrated a general increase in fibrosis-related genes in WT obese as compared to WT lean mice, with these changes attenuated in SGK1 DN obese mice (Figure 5C). Activity – and ratio-of matrix metalloproteinases (MMP) and tissue inhibitors of matrix metalloproteinases (TIMP) is known to affect the maintenance and turnover of the extra-cellular matrix (ECM).^37–39^ High fat diet feeding was associated with a significantly increased expression of tissue inhibitor of metalloproteinase 1 (TIMP1) and plasminogen activator inhibitor 1 (PAI-1), both of which are associated with increased ECM turnover. However, the ratio of TIMP1/MMP2 was significantly increased in obese WT atria as compared to lean WT mouse atria, with this difference minimized in the obese SGK1 DN atria (Figure 5D). A similar trend was noted in the ratio of TIMP1/MMP9. Overall, these data demonstrate attenuation of obesity-related profibrotic signaling in SGK1 DN atria, consistent with the previously described finding of reduced ventricular fibrosis in the TAC model.

**Figure 5:**
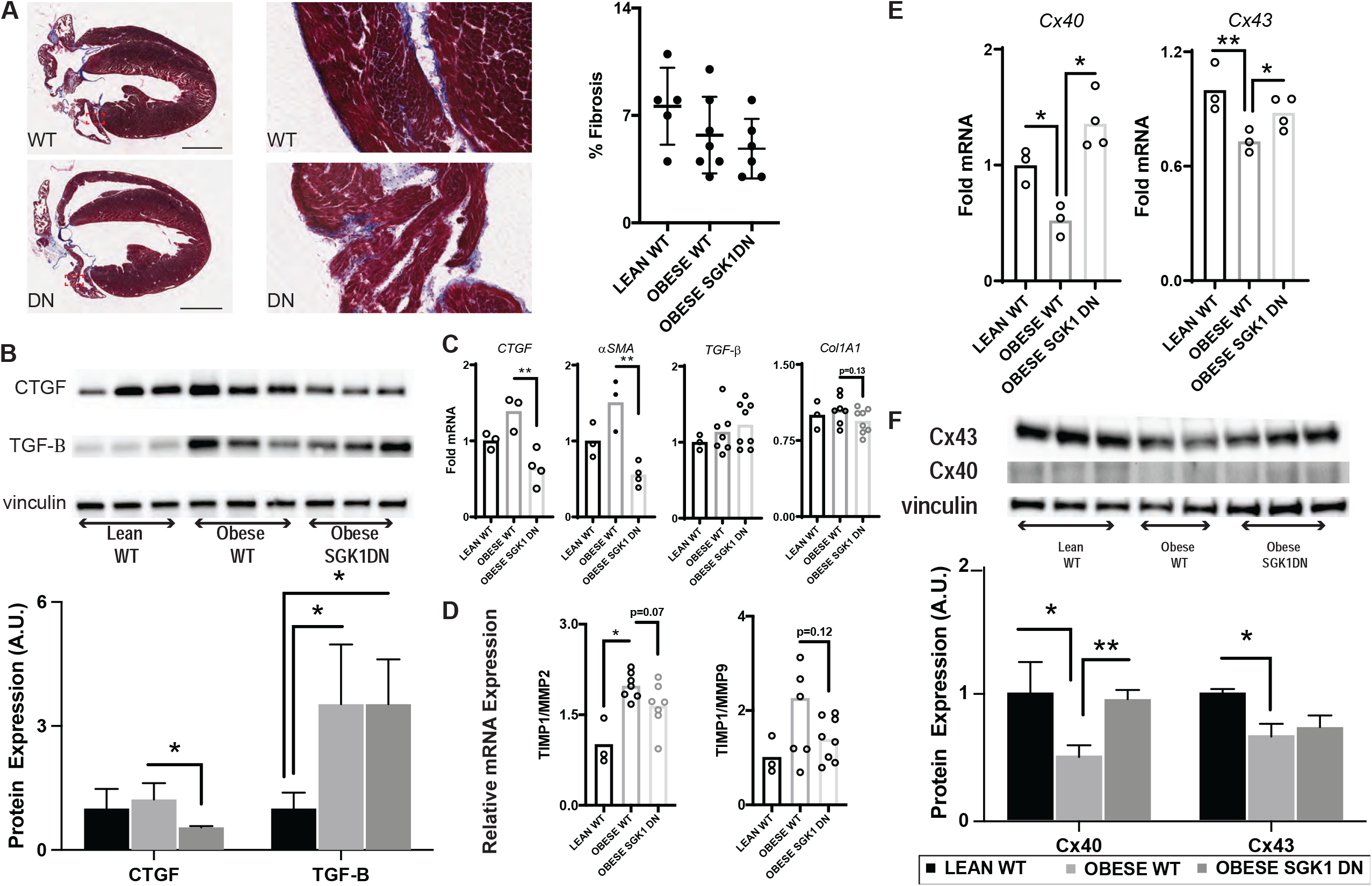
SGK1 genetic inhibition prevents obesity-induced atrial fibrotic signaling. *A*, Example Masson Trichrome stained tissue sections from mouse atria with quantification of fibrosis shown to the right. Images to the right are zoomed into the region delineated by the red dotted box. The scale bar = 2 mm. *B*, Atrial CTGF and TGF-β protein expression. *C*, Expression of fibrosis-related genes in lean and obese atria. *D*, Ratio of matrix metalloproteinase mRNA transcripts in atrial tissue. *E*, Expression of connexin mRNA transcripts. *F*, Connexin protein expression measured by Western Blot, with quantification below.

Multiple animal models have demonstrated the obesity related downregulation of cardiac connexin proteins.^11,12^ We confirmed these results in our study by determining the level of gene and protein expression. Quantitative PCR showed a decrease in atrial expression of both connexin 40 and 43 in obese as compared to lean mice, but the decrease was mitigated or reversed by SGK1 inhibition. These findings were extended to protein translation, as immunoblotting revealed obesity-related decrease in atrial protein expression of connexin 40 and 43 which were ameliorated with SGK1 inhibition. The data presented here suggest a protective effect of SGK1 inhibition in terms of pro-fibrotic signaling and cell-cell connectivity. (Figure 5E-F)

### Genetic inhibition of SGK1 prevents obesity-induced inflammation and stress signaling

Inflammation is thought to be involved in obesity induced pathology (including fibrosis),^40^ and we suspected that SGK1 signaling may contribute to pro-inflammatory pathways in the heart. We therefore evaluated the effect of SGK1 inhibition on atrial inflammation caused by obesity with quantitative PCR of inflammatory genes in WT lean, WT obese, and SGK1 DN obese mice. The genes of interest were either cytokines known to be involved in obesity-related inflammation or those thought to lie downstream of either mineralocorticoid or SGK1 signaling.^32,41–44^ Once again, cardiomyocyte specific genetic inhibition of SGK1 minimized obesity induced increases in genes related to inflammation. (Figure 6A) Given that SGK1 inhibition is restricted to cardiomyocytes, it is not surprising that plasma levels of circulating cytokines such as IL-6 and CRP were not impacted. (Supplemental Figure 3)

**Figure 6:**
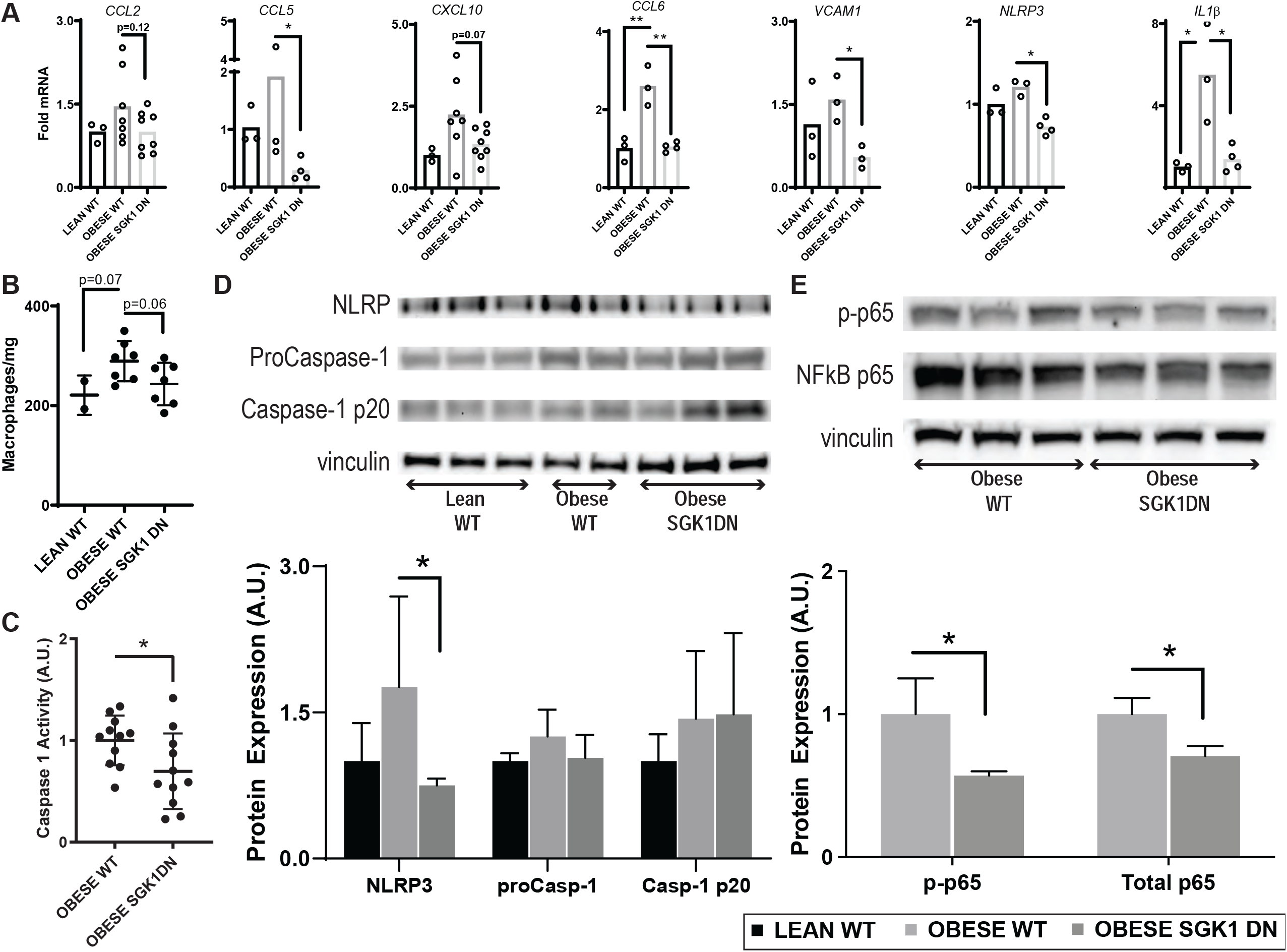
SGK1 genetic inhibition prevents obesity-induced atrial inflammatory signaling. *A*, Atrial inflammatory gene transcript levels. *B*, Atrial macrophage content in lean and obese atria as measured by flow cytometry. *C*, Caspase 1 activity in obese atrial tissue as measured with a commercially available assay. *D*, Western blots of atrial NLRP3 and caspase subunits with quantification below. *E*, Western blots of atrial NFΚB subunit p65 and its phosphorylated isoform with quantification below.

Of note, there was a marked obesity-induced atrial IL-1β expression, which was abolished with genetic inhibition of SGK1 (Figure 6A). SGK1 activation has been shown to affect expression of chemokine ligand 2 (CCL2), which is involved in the recruitment of inflammatory cells such as macrophages.^41^ Although there was only a modest difference in the expression of the CCL2 gene between the WT and SGK1 DN mouse atria, we interrogated this pathway further with flow cytometry and determined that SGK1 genetic inhibition abrogated a trend towards an increased atrial macrophage content caused by obesity. (Figure 6B) Although one could presume sheer difference in quantity of inflammatory cells as the source of the increase in IL-1β, we pursued this avenue further by investigating downstream pathways.

We studied two pathways that we suspected may have been inhibited with SGK1 inhibition: the NLRP3 inflammasome and NFκB signaling. The NLRP3 inflammasome has been implicated in various obesity-related pathologies, including AF, and has been shown to be inhibited by pharmacologic SGK1 inhibition.^32^ Additionally, SGK1 activity has been shown to provoke nuclear translocation and hence activation of nuclear factor κB.^45–47^ We assayed inflammasome activity with a fluorometric measurement of caspase activity, which was found to be significantly lower in atrial tissue of SGK1 DN mice compared to WT. (Figure 6C) NLRP3 protein and mRNA expression both showed a trend towards an increase with HFD feeding, and SGK1 inhibition resulted in a significant decrease compared to WT littermates; there were no differences in caspase-1 protein expression (Figure 6A,6D). To further assess the role of the NLRP3 inflammasome in obesity-related AF, EP studies were performed in age-matched obese NLRP3^-/-^ mice which demonstrated an intermediate AF phenotype, with 4 out of 10 mice inducible for AF greater than one second in duration. (Table 1)

NLRP3 expression is closely linked to NFκB activation, which is mechanistically linked to SGK1 activation.^48,49^ Western blotting of the p65 subunit of NFκB revealed a significantly (p<0.05) lower protein expression of both phosphorylated- and hence transactivated form-^50,51^ and total NFκB p65 in SGK1 DN mouse atria. Overall, these data suggest that the protective effect of SGK1 inhibition in IL-1β expression may be mediated in part, by its role in modulating the NLRP3 and NFκB pathways.

### Constitutive SGK1 Activation Is Not Sufficient to Increase AF Susceptibility

To determine whether SGK1 overactivation on its own determines AF susceptibility, we performed EP studies on age matched lean SGK1 CA and WT mice. There were no differences in AF inducibility noted between the two groups. Interestingly, SGK1 CA mice demonstrated a significantly longer PR interval, and had a trend (p=0.14) towards a longer Wenckebach cycle length than their WT counterparts; these data suggest an effect on AV node function. Otherwise, there were no significant differences in EP parameters between the two lean groups.

## DISCUSSION

Obesity is a reversible risk factor for AF, and the rate of obesity continues unabated at epidemic proportions. Prior work examining the link between obesity and AF has implicated fibrosis, connexin dysregulation, inflammation, and ion channel alterations in arrhythmogenesis. Here we demonstrate that diet induced obesity upregulates atrial SGK1 transcription and activation. SGK1 inhibition is associated with a reduction in AF inducibility in a mouse model of high fat diet induced obesity. This protective effect was associated with alterations in atrial electrophysiology as well as an attenuation in the obesity-related structural remodeling and inflammation. Overall, SGK1 inhibition prevents some of the pathologic effects of obesity, making it a necessary signaling pathway in this AF model.

While cardiac SGK1 activity may be protective in acute pressure overload,^29^ persistent activation during chronic pressure overload has been shown to maladaptive.^30^ SGK1 is transcriptionally upregulated by circulating factors including glucocorticoids, mineralocorticoids, and insulin.^52^ Given the latter, we suspected that obese mice, which are known to have glucose intolerance,^53^ may have increased SGK1 expression. We demonstrate here that in fact diet induced obesity does increase SGK1 expression in the heart. Although this is a novel finding in the heart, similar models of obesity have been shown to increase SGK1 expression in the aorta,^25^ adipose tissue,^26^ and in the hippocampus.^27^

Obesity has been implicated in the development of both the triggers and substrate that underlie cardiac arrhythmias.^6^ There is discordant data regarding the effect of obesity on the cardiac sodium current. ^36,54–57^ A particularly relevant recent study of mice with diet-induced obesity, for example, demonstrated decreased expression of the sodium channel subunit Na_V_1.5 associated with a concordant decrease in *I*_*Na*_ as well as APD shortening and decreased dV/dt_max_ in obese-as compared to lean-mouse left atria.^58^ These data would seem to stand in contrast to our findings, wherein SGK inhibition in obese mice was associated with a decrease in *I*_*Na*_ but also a protective effect in terms of AF inducibility. However, despite the apparent contradiction, the same mouse model had 25% reduction in AF burden when treated with flecainide, an *I*_*Na*_ blocker.^59^ In the context of available literature, our results suggest that *I*_*Na*_ inhibition may be protective in obesity-related AF, but this incompletely explains the benefit derived from SGK1 genetic inhibition.

There is substantial data suggesting a role for obesity-induced structural remodeling in the pathogenesis of atrial fibrillation. Proposed mechanisms include interstitial fibrosis,^7,8,10^ connexin dysregulation,^11,12^ and even fatty infiltration.^7^ SGK1 activity in particular is associated with a number of fibrosing conditions throughout the body, and may be stimulated by TGF-β activity.^60^ Although the data presented here did not demonstrate macroscopic histology-detected fibrosis, we do demonstrate an increased expression of several pro-fibrotic factors, including CTGF and α-SMA, which were mitigated by SGK1 genetic inhibition. These findings further substantiate a role for SGK1 signaling in obesity-related fibrosis.^30^ MMPs and TIMPs have generally antagonistic functions and work in concert to maintain the homeostatic balance of the extracellular matrix.^37,38^ The relative changes seen here with obesity-which mimic those seen in aged, frail mouse atria-^39^ may suggest a profibrotic transcriptional programming. The protective effects of SGK1 genetic inhibition are not entirely unexpected, as they somewhat replicate the effects of cardiomyocyte mineralocorticoid receptor knockout.^61^ The role of SGK1 signaling on cardiomyocyte connexin expression are not described in literature, but there is extensive data regarding the effects of fibrosis and inflammation on connexin expression, so this may simply be a more downstream finding.

SGK1 activation has been linked to inflammation via both the NLRP3 inflammasome as well as NF-κB signaling. With respect to the latter, SGK1 directly phosphorylates IKKα, resulting in increased NF-κB activity.^49^ SGK1 dependent NF-κB activation has been demonstrated in renal collecting ducts^62^ and aortic vascular smooth muscle cells.^63^ Recent literature has posited cardiomyocyte NLRP3 overexpression as a sufficient trigger for AF,^14^ and our data regarding atrial NLRP3 expression is compelling in this context. Prior studies have demonstrated an association between SGK1 and inflammasome activity, but the exact mechanism has not been clearly delineated. In a model of hypoxia-induced pulmonary arterial hypertension, SGK1 activity was shown to be associated with hypoxic pulmonary macrophage infiltration, whereby knockout of SGK1 reduced macrophage content.^64^ Meanwhile, a specific SGK1 inhibitor was shown to reduce NLRP3 inflammasome expression and mitigate angiotensin II induced cardiac inflammation and fibrosis.^32^ In our model, obesity resulted in a marked elevation in IL-1β-a cytokine frequently cited as pro-arrhythmic-which was prevented by SGK1 inhibition, perhaps through its effects on the NF-κB and NLRP3 axes.

An interesting aspect of this study to take note of – as in the *Das et al*^30^ study regarding TAC-induced cardiac remodeling-is that our model of SGK1 knockdown is restricted to cardiomyocytes via the αMHC promoter. Yet, the effects seemingly extend to functions classically thought to be driven by non-cardiomyocytes; ECM maintenance is generally attributed to activated fibroblasts, and inflammation to monocytes. However, *in vitro*, cardiomyocytes have been shown to be capable of expressing fibrosis related transcripts,^65^ and even do so in response to leptin exposure.^66^ Murine *in vivo* models of cardiac fibrosis due to pressure overload have demonstrated an essential role for cardiomyocyte-dependent TGF-β signaling.^67^ There is limited data regarding the role of cardiomyocyte derived signaling in inflammation, but cardiomyocyte calmodulin kinase - through both NLRP3 and NF-κB – is essential to pressure overload induced cardiac inflammation.^68,69^ A particularly relevant study from *Rickard et al*^61^ demonstrated cardiomyocyte mineralocorticoid receptors as essential in mineralocorticoid and salt induced cardiac fibrosis and inflammation. This is of critical interest in the context of our data, as the SGK1 axis lies downstream of mineralocorticoid receptor signaling via transcriptional regulation.

We acknowledge several limitations in our study. At first glance, it would seem that the stark differences in AF inducibility caused by SGK1 inhibition is incompletely explained by the less pronounced mechanistic differences. We propose that SGK1 inhibition-through its pleiotropic effects-decreases AF susceptibility via multiple contributory mechanisms. Another limitation innate to any rodent studies of AF is that a “positive” finding typically represents very brief (on the order of seconds) episodes of atrial tachyarrhythmia that are induced *in vivo*; i.e. these episodes are neither spontaneous nor sustained. We would propose that generally speaking mouse models of AF are models of the substrate, with the trigger provided by pacing/stimulation.

In summary, this study suggests that pharmacologic inhibition of SGK1 should be further investigated for its therapeutic potential in obesity-related AF. In addition, this study, like others focused on obesity-related AF, describes a number of paths that may lead from obesity to AF beyond ion channel alterations. Notably, SGK1 activity seems to be required for many of these pro-arrhythmic pathways, suggesting a central role for this kinase in obesity-related AF, and thus making inhibition a potentially attractive target for intervention. Future work at targeting some of these paths may provide avenues to novel drug therapy for the increasingly prevalent obesity-related AF.

## FUNDING SOURCES

Dr. Aneesh Bapat was supported by the NIH grant T32HL007604. Dr. Ling Xiao was supported by AHA 20CDA35260081. Dr. Saumya Das was funded by AHA SFRN SFRN35120123 and AHA 18IPA34170109. Dr. Matthias Nahrendorf was supported by NIH grant R35 HL139598. Dr. Maximillian J. Schloss was supported by Deutsche Forschungsgemeinschaft (SCHL 2221/1-1).

## DISCLOSURES

SD, DM and AR are founding members of Long QT Therapeutics, which did not play any role in the funding or design of this study. M.N. has received funds or material research support from Alnylam, Biotronik, CSL Behring, GlycoMimetics, GSK, Medtronic, Novartis and Pfizer, as well as consulting fees from Biogen, Gimv, IFM Therapeutics, Molecular Imaging, Sigilon and Verseau Therapeutics.

## SUPPLEMENTAL TABLE

**Supplemental Table 1:**
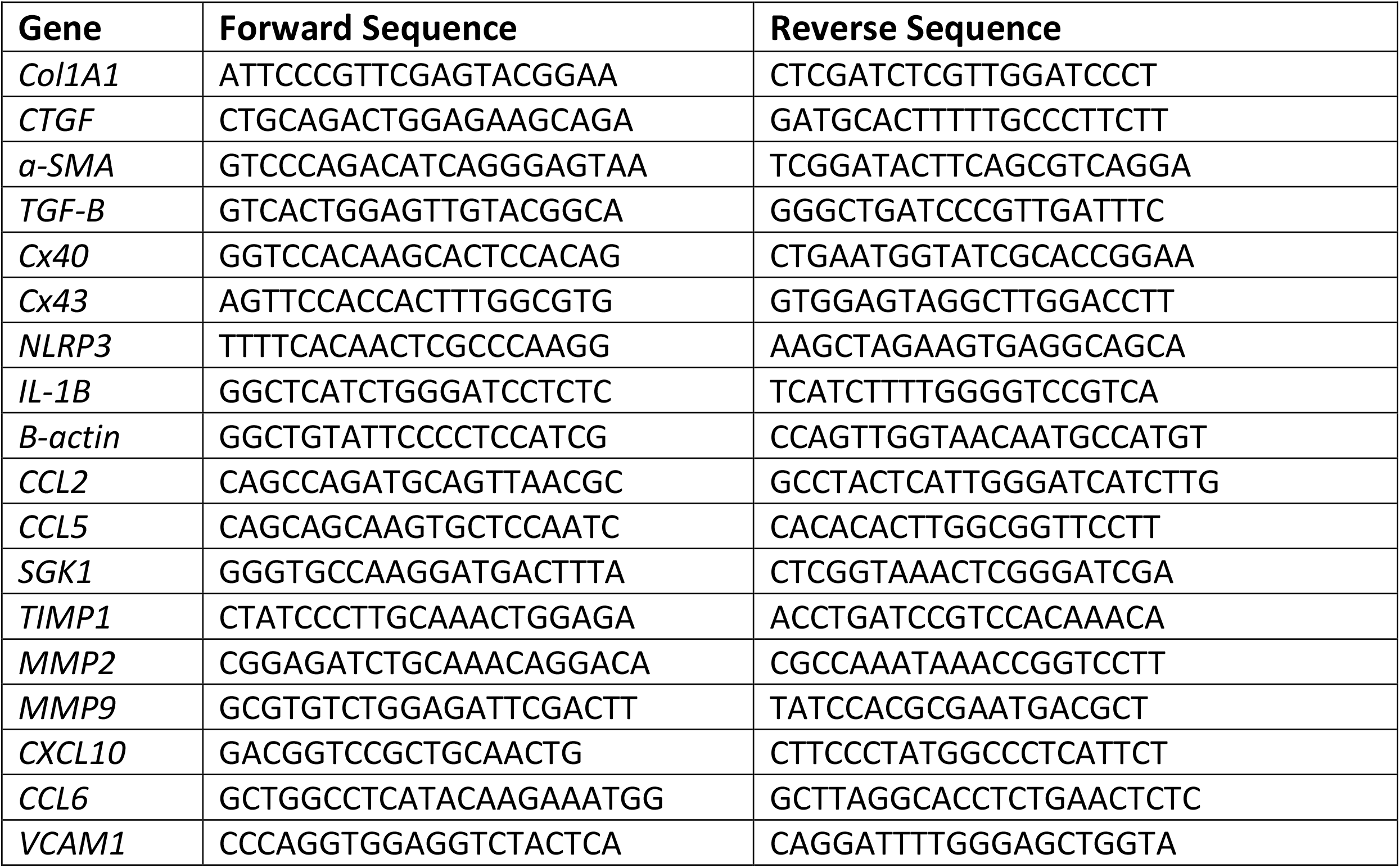
Primer sequences for genes of interest assayed by quantitative PCR.

## SUPPLEMENTAL FIGURES

**Supplemental Figure 1:**
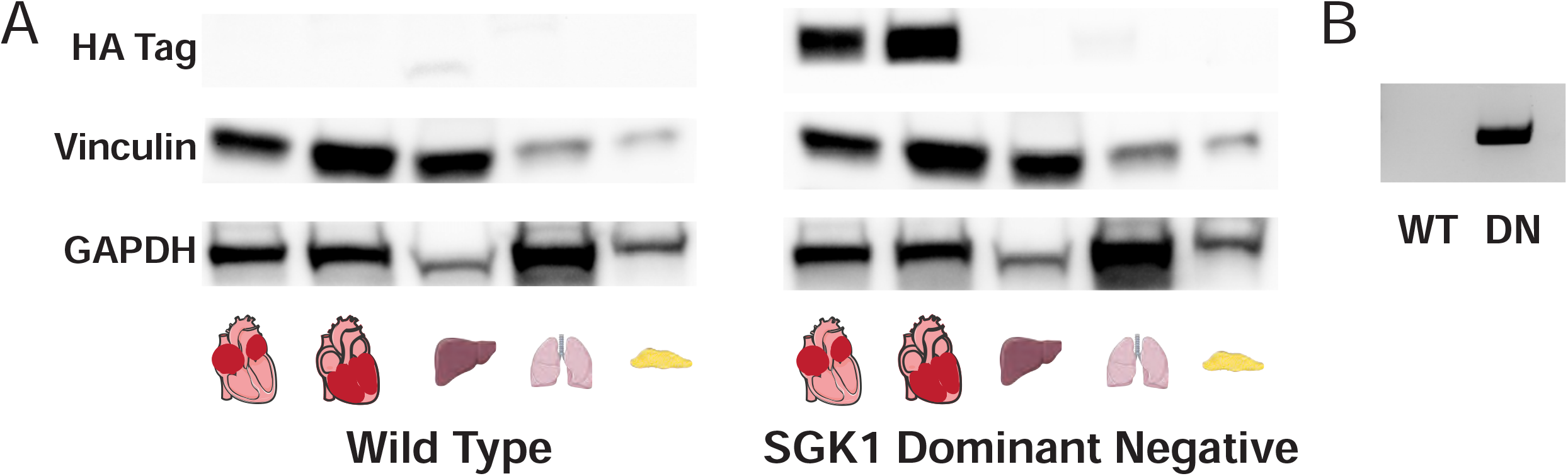
Cardiac specificity of SGK1 DN transgene. *A*, Western blots of hemagglutinin (HA) epitope tag present in only the atrial and ventricles of SGK1 DN mice, but not in non-cardiac tissues or in WT mice. *B*, Example DNA blot demonstrating presence of SGK1 transgene in SGK1 DN, but not WT, mice.

**Supplemental Figure 2:**
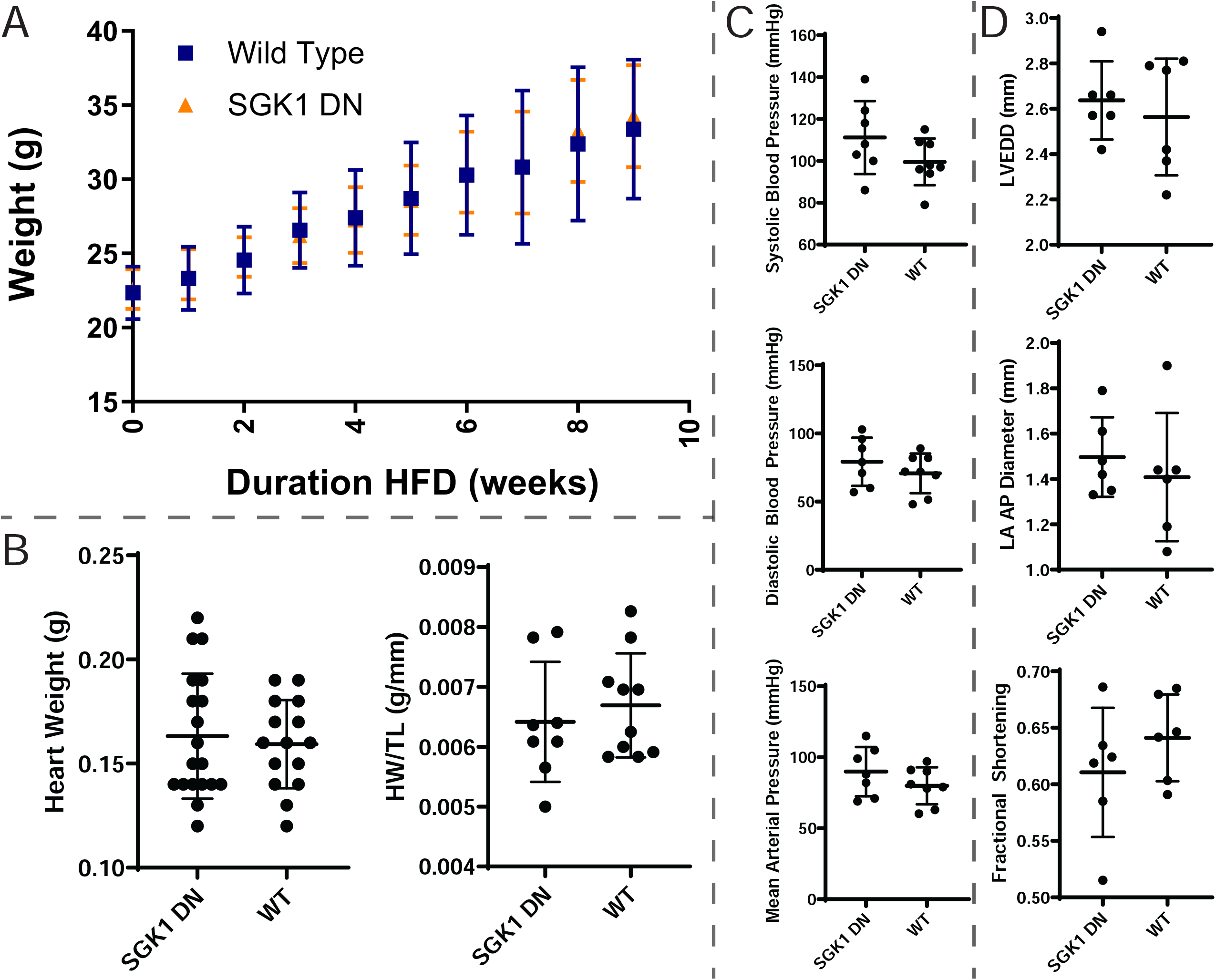
Phenotyping HFD-fed WT and SGK1 DN mice. *A*, Mouse body weight during HFD feeding. *B*, Heart weight and heart weight indexed to tibial length in mice. *C*, Blood pressure parameters measured after HFD feeding. *D*, Transthoracic echocardiography measured cardiac structural/functional parameters after HFD feeding.

**Supplemental Figure 3:**
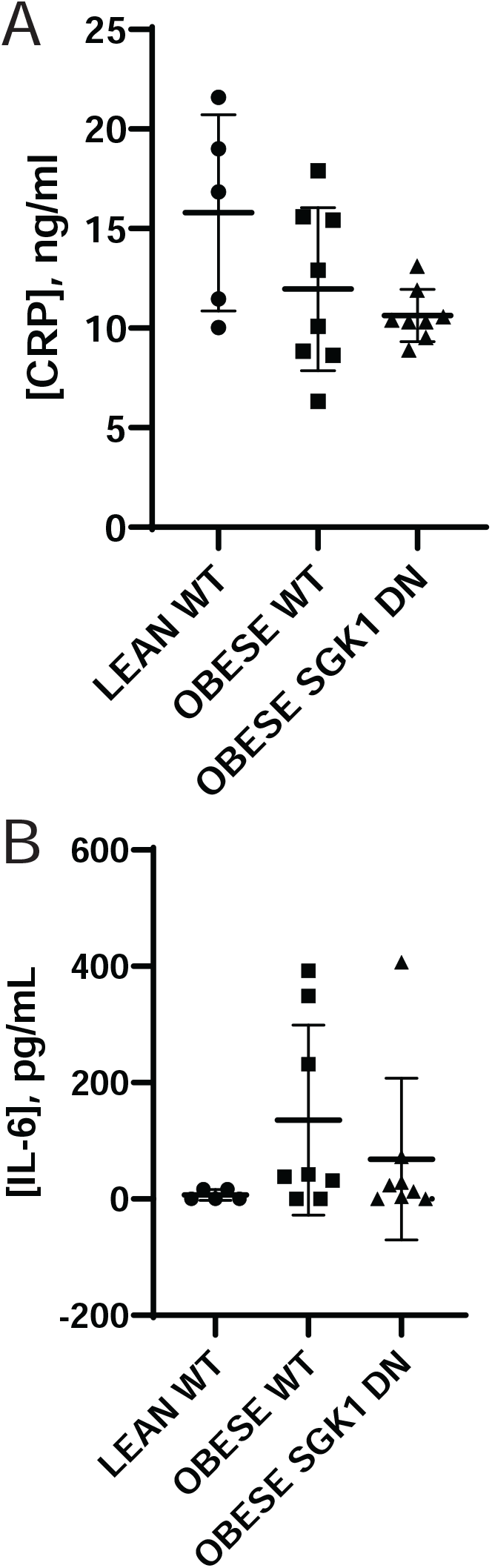
Plasma markers of systemic inflammation. *A*, Plasma levels of C-reactive protein (CRP). *B*, Plasma levels of interleukin 6 (IL-6).

## Notes

### Competing Interest Statement

The authors have declared no competing interest.

